# Human activity is altering the world’s zoogeographical regions

**DOI:** 10.1101/287300

**Authors:** Rubén Bernardo-Madrid, Joaquín Calatayud, Manuela González-Suarez, Martin Rosvall, Pablo M. Lucas, Marta Rueda, Alexandre Antonelli, Eloy Revilla

**Affiliations:** Department of Conservation Biology, Estación Biológica de Doñana (EBD-CSIC), Sevilla, Spain; Department of Life Science, Universidad de Alcalá, Alcalá de Henares, Spain; Deparment of Biogeography and Global Change, Museo Nacional de Ciencias Naturales (MNCN-CSIC), Madrid, Spain; Integrated Science Lab, Department of Physics, Umeå University, 901 87 Umeå, Sweden; Ecology and Evolutionary Biology, School of Biological Sciences, University of Reading, Reading, UK; Department of Wildlife Conservation, Institute of Nature Conservation (IOP-PAS), Kraków, Poland; Gothenburg Global Biodiversity Centre, Box 461, SE-405 30 Göteborg, Sweden; Department of Biological and Environmental Sciences, University of Gothenburg, Box 461, 405 30 Göteborg, Sweden; Gothenburg Botanical Garden, Carl Skottsbergs gata 22A, SE-41319 Göteborg, Sweden; Department of Organismic and Evolutionary Biology, Harvard University, 26 Oxford St., Cambridge, MA 02138 USA

**Keywords:** Conservation, Invasion biology, Extinction, Vertebrates

## Abstract

Human activity leading to both species introductions and extinctions is widely known to influence diversity patterns on local and regional scales. Yet, it is largely unknown whether the intensity of this activity is enough to affect the configuration of biodiversity at broader levels of spatial organization. Zoogeographical regions, or zooregions, are surfaces of the Earth defined by characteristic pools of species, which reflect ecological, historical, and evolutionary processes acting over millions of years. Consequently, it is widely assumed that zooregions are robust and unlikely to change on a human timescale. Here, however, we show that human-mediated introductions and extinctions can indeed reconfigure the currently recognized zooregions of amphibians, mammals, and birds. In particular, introductions homogenize the African and Eurasian zooregions in mammals; reshape boundaries with the reallocation of Oceania to the New World zooregion in amphibians; and divide bird zooregions by increasing biotic heterogeneity. Furthermore, the combined effect of amphibian introductions and extinctions has the potential to divide two zooregions largely representing the Old and the New World. Interestingly, the robustness of zooregions against changes in species composition may largely explain such zoogeographical changes. Altogether, our results demonstrate that human activities can erode the higher-level organization of biodiversity formed over millions of years. Comparable reconfigurations have previously been detectable in Earth’s history only after glaciations and mass extinction events, highlighting the profound and far-reaching impact of ongoing human activity and the need to protect the uniqueness of biotic assemblages from the effects of future species introductions and extinctions.

## Main text

Zoogeographical regions, or zooregions, are areas of the Earth defined by a characteristic pool of species, and are the outcome of long (co)evolutionary histories, local and global extinctions, and colonization events. The significance of diversification (i.e., the balance between speciation and extinction) and colonization processes in generating zooregions varies in time and space because of multiple factors acting on different scales, such as tectonic movements, climate change, geographic barriers, and ecological interactions (1, 2). Because these processes have been acting for millions of years, zooregions are often assumed to be resilient on a human timescale. However, the current spread and magnitude of human activity may challenge this assumption (3).

Previous studies have shown how human activity is affecting the dynamics of diversification and dispersal, triggering both local and global extinctions by reducing and fragmenting the habitat of species (4), by direct persecution (5, 6), and indirectly through extinction cascades (7). Human activity is also altering the dynamics of dispersal by breaking down natural barriers for some species through human trade and travel (8), while limiting large-scale movements for others (9). These processes have already produced changes in the diversity patterns of many species on both local and regional scales (10, 11); however, whether the effects of human activity scale up to the highest organizational levels of life on Earth remains largely unknown.

Theoretically, introductions and extinctions may trigger three types of zoogeographical changes (Fig. 1). Species introductions can homogenize zooregions that encompass native and introduced ranges (3, 12), but can also differentiate the invaded and non-invaded areas possibly leading to division (13). Complementarily, extinctions may cause homogenization if species unique to a zooregion go extinct (12, 14, 15), but may also cause division if widespread species go extinct, increasing regional heterogeneity (16, 17). Finally, depending on the final distribution of species, introductions and extinctions may redefine the boundaries of zooregions without varying the overall number. For instance, an area may be reassigned to a neighboring zooregion if species characteristic of that region invades it, or if the area loses some of the species that characterized it. In summary, introductions and extinctions have the theoretical potential to homogenize, divide, and redefine zooregions.

**Fig. 1.**
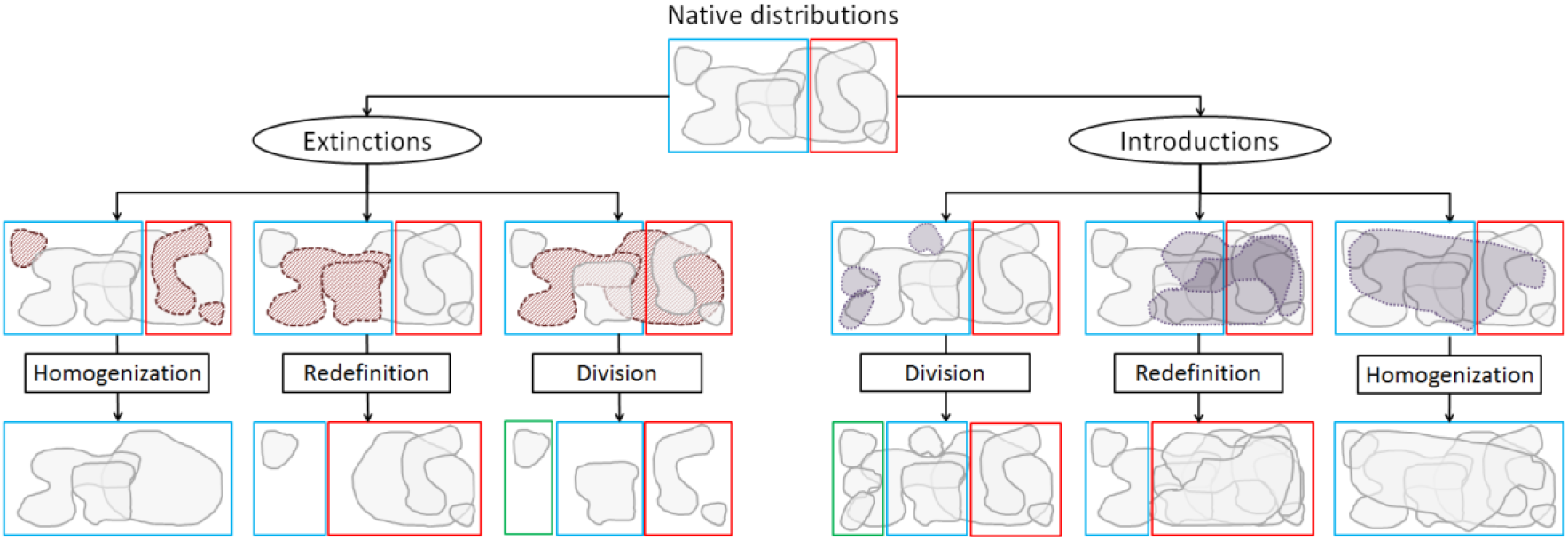
Theoretical changes in the delineation of zooregions as a consequence of human-mediated extinctions and introductions of species. The blue, red, and green rectangles represent three hypothetical zooregions with inside polygons reflecting hypothetical species distributions. The diagram has three levels: (i) at the top of the figure there is a natural (pre-human impact) scenario with two zooregions (blue and red); (ii) this natural state is disrupted by either extinctions (left; species disappearing are marked by red polygons with dashed lines) or introductions (right; introduced species are marked by gray polygons with solid lines); (iii) the third level (human-affected) represents the outcome of these extinctions and introductions, and their changes to the original zooregions. The actual observed changes depend on how the composition of the species is affected.

Current efforts to map the native and introduced distribution of species, together with the development of robust analytical techniques to delineate zooregions, allow the evaluation of whether and how human activity modifies zooregions. Indeed, detailed information about the native and introduced distribution ranges of species can help to determine whether human-mediated introductions are currently affecting zooregions. In addition, the information on species extinction risk may be used to shed light on potential effects of species losses by simulating extinctions of the most threatened species. Finally, the use of these data along with recently developed classification techniques provides a unique opportunity to explore the potential causes of observed zoogeographical changes. In particular, network theory has provided powerful tools to define biogeographic regions (17–22) and identify the set of species that characterize each zooregion (18, 19). Importantly, these network tools also allow identifying the role of particular species within each zooregion (19, 23), making it possible to measure the robustness of zooregions against changes in species composition and to quantify the impact of introductions and extinctions on zooregion delimitation.

Here we assessed how the observed introductions and the potential extinction of the most threatened species could affect the configuration of zooregions in amphibians (6,103 species), non-marine mammals (5,001 species), and non-pelagic birds (9,834 species). First, we used a network community detection algorithm (20, 21, 24) to delineate the zooregions of four scenarios: (i) native, (ii) introduced, (iii) extinct, and (iv) the combination of introduced plus extinct (see Material and Methods). Second, we quantified differences between the scenarios in the configuration of the zooregions. Finally, to provide a deeper understanding of the zoogeographical changes, we measured the robustness of zooregions and located and quantified the impact of introductions and extinctions. Our results showcase that human-mediated introductions are already substantially affecting the world’s zooregions and that this impact could escalate if threatened species go extinct. Interestingly, our findings also highlight that the robustness of zooregions largely explain zoogeographical changes.

## Results

### Native zooregions

We used species native distributions to delineate natural zooregions hierarchically (20, 21), following the standard approach in biogeographical studies (25, 26). As expected, our native zooregions are similar to the ones first proposed by Wallace (1). However, for amphibians and birds, we detect an additional higher hierarchical level, delineating two main zooregions corresponding to the New and the Old World (Fig. 2, A1 and B1). Interestingly, our highest and lowest hierarchical levels (hereafter major and minor zooregions) are similar to the ‘realms’ and ‘regions’ obtained with other methodological approaches (20, 25–27). These are the most commonly used spatial units in biogeography (28, 29), and, therefore, we used the major and minor zooregions of native scenarios as a baseline for assessing whether human-mediated introductions and extinctions affect the configuration of zooregions.

**Fig. 2.**
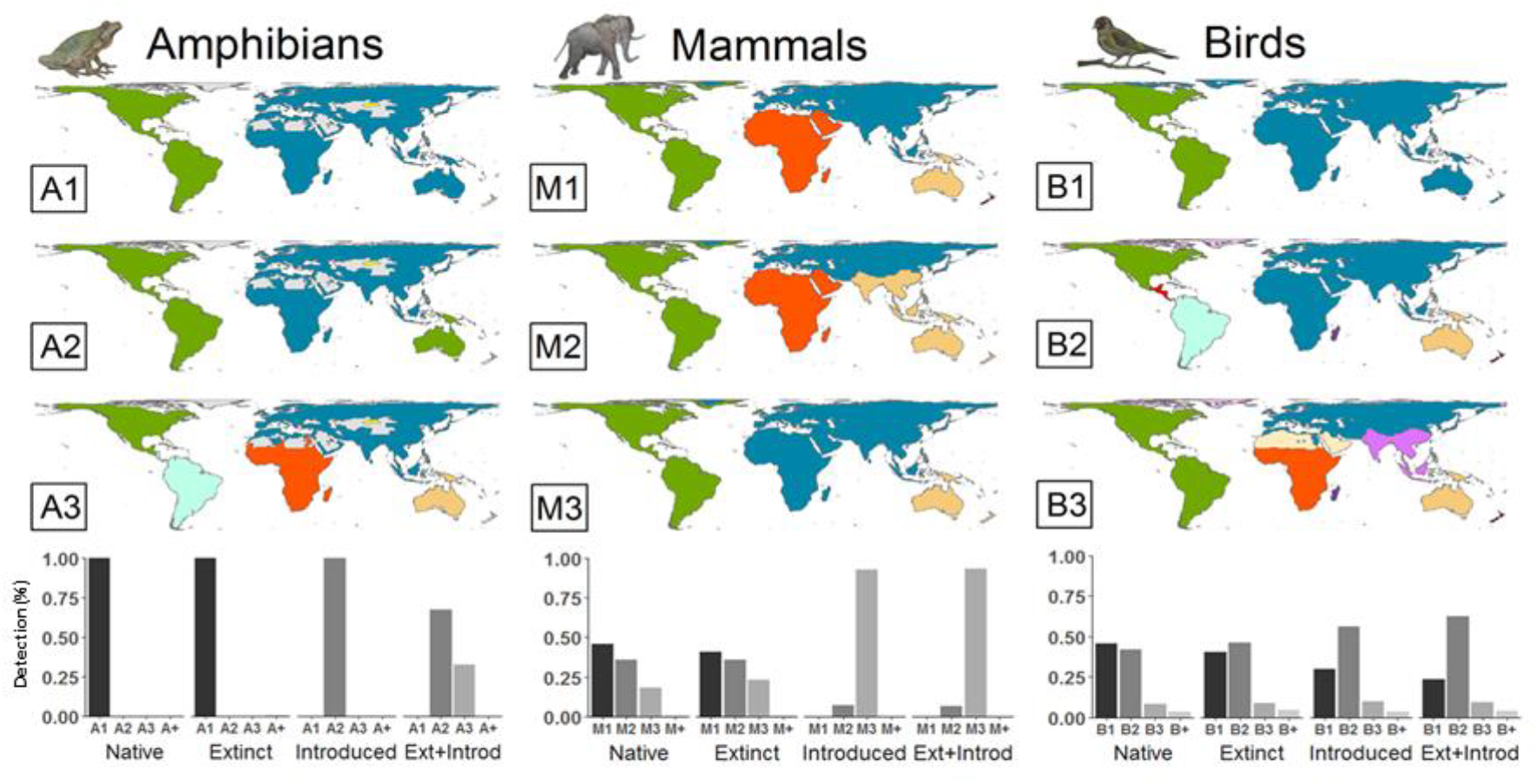
Observed introductions and potential extinctions increase the probability of detecting notably modified zooregions in amphibians, mammals, and birds. Maps with the zoogeographical delineations of the three most probable solutions for the major zooregions of amphibians, mammals, and birds (labeled as A1–A3 for amphibians, M1–M3 for mammals and B1–B3 for birds) and histograms with their probability of detection. In amphibian delineations, gray areas are those where amphibians are not present according to the IUCN. The histograms also show the joint probability of detection of less frequently observed solutions, not shown in maps (see SI Appendix, Table S1 and Fig. S1-3 for details).

### Introduced scenario

We delineated our introduced scenario using the same methodology as in the native one but also including the introduced ranges of species. To robustly compare scenarios, we considered the uncertainty inherent in delineating zooregions. Because of areas occupied by many characteristic species from diverse regions (e.g., transition areas) may be assigned to diverse zooregions, a scenario may be delineated with alternative similarly supported solutions. Consequently, for each scenario, we identified the alternative supported solutions and calculated their probabilities to be detected (see Material and Methods for details). We then measured how the detection probability of existing solutions differs between scenarios using Schoener’s index (hereafter D) (30). If introductions (or extinctions, discussed below) affect the configuration of zooregions, we expect low similarities in the probability of detection of the solutions in different scenarios (with D approaching 0).

In amphibians, our results show a single solution in the introduced scenario that differs from the native one (D = 0, Fig.2, A2). In the new solution, the major zooregions representing the New and the Old World are redefined due to the relocation of the Australian region into the New World zooregion. In mammals, our results also show important changes in the detection probability of solutions across scenarios (D = 0.25, Fig.2). Specifically, mammal introductions increase the probability of detecting a homogenized major zooregion including Africa, Eurasia, and the Indo-Malayan regions (Fig.2, M3). Additionally, our results show a smaller effect in birds (D = 0.83, Fig.2). Nevertheless, bird introductions decrease the detectability of the two major zooregions corresponding to the New and the Old World (Fig.2, B2). Regarding minor zooregions, mammalian introductions increase the similarity between the North African and Arabian minor regions (D = 0.38; SI Appendix, Fig. S4, m2), but we found no effects in amphibians or birds (D = 1 and D = 0.99, respectively; SI Appendix, Fig. S4). Overall, our results reveal that introductions trigger redefinition, homogenization, and division of the major zooregions of amphibians, mammals, and birds respectively, as well as homogenization of the minor zooregions of mammals.

### Extinct scenario

In addition to the effects of current introductions, zooregions could also be affected by human-mediated extinctions. We considered this possibility by simulating a hypothetical scenario in which the most threatened species, those listed as Critically Endangered and Endangered by the International Union for Conservation of Nature (IUCN) Red List, become globally extinct. As many threatened species are endemic to one zooregion, we expect a homogenization effect (Fig.1). However, independently of the vertebrate group and spatial scale (major and minor zooregions), we found no relevant effects on zooregions (SI Appendix, Table S2).

### Introduced and extinct scenario

Because introductions and extinctions happen in concert, we also considered a hypothetical combined scenario. We delineated this scenario using the same methodology as in the introduced scenario but also simulating the extinction of the most threatened species. In general, detected changes are similar to those in the introduced scenario (SI Appendix, Table S2), and support the apparent limited effects of tested extinctions found in the extinct scenario. Nonetheless, the combination of amphibian introductions and extinctions triggers new changes in which zooregions representing the New and the Old World are no longer detected (Fig.2, A3).

### Robustness and impacts

We explored to what extent the robustness of native zooregions and the location and magnitude of the impacts of introductions and extinctions explain zoogeographical changes. We quantified these effects by using the information about the characteristic species —species mainly distributed in that zooregion— and non-characteristic species —overlapping but not characteristic species— in each grid cell of the zooregions. To measure robustness, we quantified (i) how each characteristic species represent its zooregion by an indicator value (31), (ii) how much each non-characteristic species reduces the uniqueness of the zooregion by a diffusion value, and (iii) the overall richness of characteristic and non-characteristic species. For instance, areas with many characteristic species and few non-characteristic ones would be highly robust. To quantify impact, we calculated to what extent the introductions and extinctions reduce the indicative value and increase the diffusion one (see Material and Methods for details). For instance, areas that lose several characteristic species and gain non-characteristic ones would be highly impacted.

Our results show that highly impacted zooregions partially coincide with zoogeographical changes. For example, they coincide for the relocation of the Australian and South American region in amphibians, the homogenization of Africa and Eurasia in mammals, and the differentiation of the Americas in birds (Fig.3B). On the other hand, some areas with high impacts, such as the American region in mammals or the African and Eurasian regions in birds, show no changes. Interestingly, differences in the robustness of zooregions may explain why similar impacts do no trigger changes in those areas (Fig.3A). For instance, variations in robustness may explain why mammals and birds experience smaller changes in their detection probabilities compared with amphibians. Moreover, differences in robustness between major and minor zooregions also agree with the magnitude of observed changes (Fig.3A and see Appendix Fig.S8). Thus, while there is no change without impact, robustness stands out as the single best descriptor of zoogeographical changes.

**Fig. 3.**
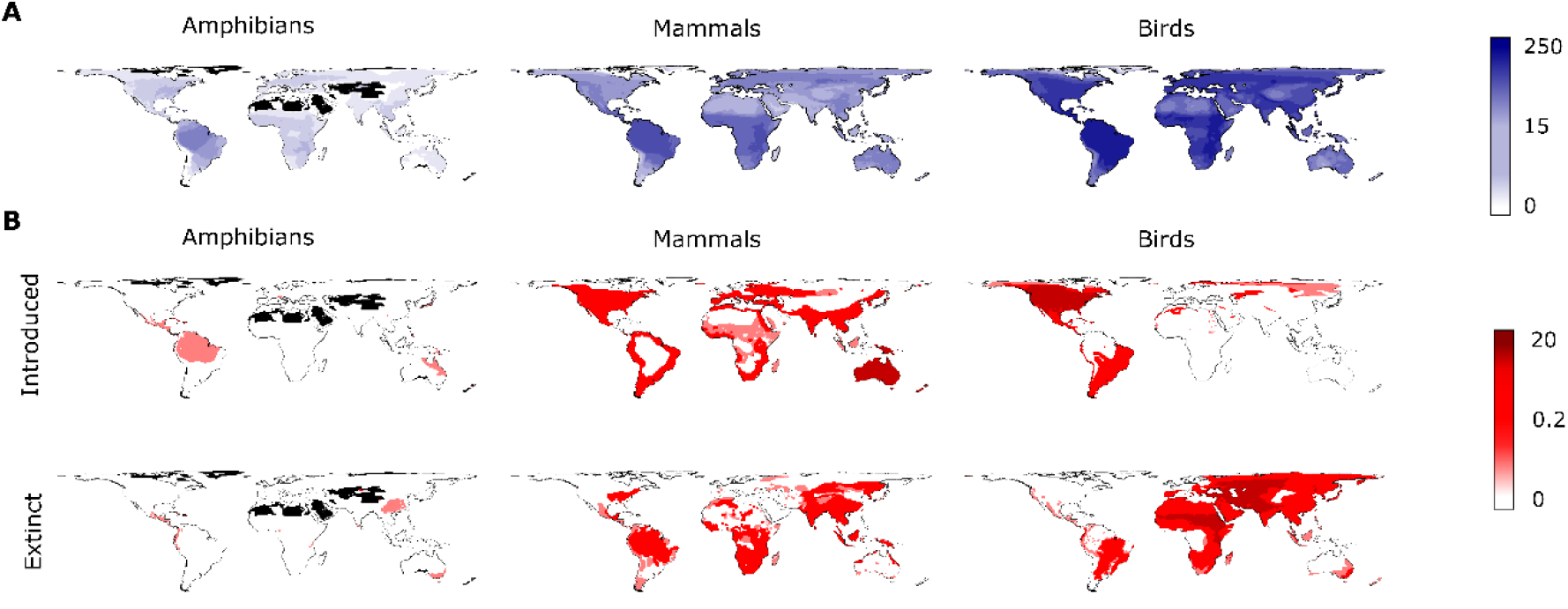
Maps representing robustness and impact under different scenarios, showing how both vary across regions and vertebrate groups. We represent the (A) robustness and (B) impact values for the grid cells of the major zooregions, calculated using the most frequent solution for each vertebrate group. (A) Light and dark blue colors indicate low and high relative robustness, respectively. (B) Light and dark red colors indicate low and high relative impact, respectively.

## Discussion

Our results demonstrate that species introductions are already modifying the world’s zooregions of amphibians, mammals, and birds at different spatial scales across the Earth’s surface. Contrary to general assumptions, human activity is not only triggering homogenizations on a global scale but also other effects such as divisions and redefinitions. Importantly, how impacts lead to zoogeographical changes is a complex issue and is affected by the intrinsic properties of the zooregion such as robustness, revealing the value of going beyond delineations for more effective biodiversity conservation.

Human-mediated introductions and extinctions of species are heterogeneously affecting zooregions across vertebrate groups and spatial scales. Introductions homogenize the African and Eurasian major and minor mammalian zooregions by making their already similar faunistic ensembles even more similar (32, 33), redefine one of the least robust areas of the major amphibian zooregions (Fig.3A), and increase the internal heterogeneity of the bird major zooregions due to intercontinental introductions (34). Although potential extinctions appear to have little effects on zooregion delimitations when considered alone, the combined effects of current introductions and extinctions are likely to trigger important modifications, as shown for the major zooregions of amphibians. Together, these findings shed light on the important consequences of human activity on the largest biodiversity units.

It should be noted that our scenarios are likely underestimating the influence of humans on zooregions. First, the introductions recorded by the IUCN may not be exhaustive, especially considering the difficulties and time-lags in tracking ongoing introductions (35). Second, the number of current species and their assumed native range already reflect past human actions (e.g., Holocene megafaunal extinctions, (36, 37)). Additionally, the current native distribution may not reflect recent range contractions due to habitat loss and climate change (38). Thus, despite the substantial changes reported, our findings are in fact conservative, even when simulating potential extinctions. Current efforts are being made to develop accurate databases including this information. When these data are available we may detect even more severe impacts.

The intrinsic properties of zooregions —i.e., robustness— and the location and magnitude of human activities explain most of the changes in the configuration of zooregions. Remaining unexplained changes may be due to more complex and untraceable consequences of extinctions and introductions. For instance, changes are difficult to predict in areas where many introduced species from several zooregions occur (e.g., mammal introductions from America and Eurasia to Oceania). Robustness, however, may explain most of the changes and the differences among vertebrate groups and spatial scales. The variation in robustness may in turn be explained by the environmental and topographic conditions surrounding zooregions (2), and the idiosyncratic dispersal abilities of each group. The high dispersal capabilities of most birds allow them to occupy wide extensions of their associated zooregions, hence having higher indicative values (see Appendix, Fig.S9). Consequently, zooregions of birds have characteristic species with higher indicative values and, therefore, their zooregions are more robust than those of mammals, which in turn, are more robust than those of amphibians (the most dispersal-limited group analyzed). Similarly, the indicative value of the characteristic species may also explain differences in robustness and detectable changes between major and minor zooregions. For a characteristic species, its indicative value depends on how much of its associated zooregion it occupies. Because there are fewer species with very wide distribution ranges (e.g., continental), thus, fewer species are highly indicative in major zooregions, making them less robust than minor zooregions (see Appendix, Fig.S9). These results show that robustness is a useful metric to identify vulnerable biogeographic areas, and thus, it could be a potentially useful tool in wide-scale biodiversity conservation.

Our findings show that human activity is changing the arguably largest conservation targets for a biologically diverse world (39). A short period of human influence on the biosphere is altering biotic entities formed over millions of years of (co)evolution, as a result of the migration of lineages across the planet, speciation, extinctions, and adaptations to local and regional environmental and biotic conditions (1, 2, 25). Moreover, modifications in species composition of zooregions might bring important consequences in the evolutionary (40–43) and ecological (44, 45) processes that characterize regional biotas. To mitigate further effects of human activity and safeguard the future of life on Earth, we should not only conserve taxonomic, functional, and phylogenetic biodiversity (46) but also greatly reduce the spread of alien species into new biogeographical regions, and protect the uniqueness of species assemblages.

## Materials and Methods

### Data

We obtained species range maps from the IUCN (47) and BirdLife International (Birdlife) (48). For our analyses, we selected polygons with presence classified as “Extant” or “Probably Extant”. We converted ranges into presence/absence maps in a regular global grid with cells of 111km x 111km (18, 25). We limited the analyses to the emerged land surface of the world (identified using the country’s layer of IUCN), and in particular, to cells with ≥50% of their area classified as land following previous studies (26). After removing subspecies and species that did not overlap with our grid we ended up with 6,103 amphibian species, 5,001 mammal species, and 9,834 bird species.

To explore the effects of introductions and extinctions, we generated two actual scenarios: native – based on the current native distribution range of extant species; and introduced – based on the native and introduced ranges of extant species, describing the current species distributions. We also generated two hypothetical scenarios: extinct – derived from the native scenario but removing the most threatened species and those listed as Critically Endangered or Endangered on the IUCN Red List, thus simulating their global extinction; and introduced plus extinct – considering both introductions and extinctions simultaneously.

### Zoogeographical delineations

We delineated the zooregions for amphibians, mammals, and birds under the above-described four scenarios. For these analyses, we used network community detection techniques (19–22, 49). Current implementations in network algorithm do not allow the incorporation of phylogenetic information prior to the clustering analyses, but identify the characteristic and non-characteristic species of each zooregion (18, 49). This dual classification is required to measure the robustness and impact in zooregions (see below). First, for each scenario, we created presence-absence matrices based on the occurrence of species in grid cells. Second, we treated the presence-absence matrices as bipartite networks (18). Third, we used the community detection algorithm known as the “map equation” (20, 21, 24) to detect zooregions (see Appendix A for details). Fourth, we obtained the assignment of each grid cell to the major and minor hierarchical levels (50). To consider the uncertainty delineating zooregions, we repeated these steps 1,500 times for each scenario. We found that 1,500 analyses are enough to describe all the alternative supported solutions of each scenario (see Appendix B for details, and SI Appendix, Fig.S10).

### Comparisons of the scenari

We assumed that highly similar zoogeographical delineations represent the same solution. Therefore, we grouped the delineations by their similarity and calculated the probability of obtaining each alternative supported solution in each scenario. Finally, we studied the differences in the detection probability of those solutions across scenarios (see workflow, Fig.S11). For each vertebrate group, we calculated the similarity between each pair of the 6,000 zoogeographical delineations –1,500 per scenario. To calculate the similarity, we developed a generalized version of the Jaccard index. We calculated the Jaccard index between all pairs of zooregions 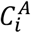, 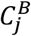 in two delineations *C*^*A*^ and *C*^*B*^ and weighted them by the fraction of all grid cells n in the intersection. This gives the weighted Jaccard partition index (equation 1)

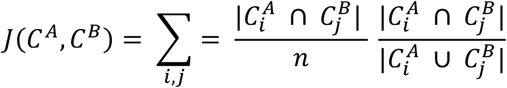

We preferred this index over others because it roughly represents the proportion of overlap between zooregions of two delineations. In other words, it approximately measures the overlapped proportion of Earth’s surface. Subsequently, we used the similarity values to group highly similar delineations, so that each group of delineations represents alternative supported solutions. To find the groups, we performed a community detection analysis by generating a unipartite network where the nodes were the delineations, and the links were the similarity values. To ensure that delineations within a cluster were very similar (i.e., at least 90% of the Earth’s surface has the same zooregions), before performing the clustering, we removed links with a similarity value below a 0.90 threshold. Once we obtained the clusters, we computed the detection probability of the solutions in each scenario as the proportion of delineations grouped in each cluster (SI Appendix, Table S1). Finally, we assessed the effects of introductions and extinctions by comparing the probability distribution of the solutions among scenarios using Schoner’s D Index (30). To ensure that our results are independent of our threshold (0.90), we repeated the analyses using different similarity threshold (0.85, 0.90, and 0.95), and we found no differences between similarity thresholds (see SI Appendix, Table S2).

### Robustness and impact indexes

To estimate the robustness of grid cells, we used to the indicator value of characteristic species and the diffusion value of non-characteristic ones present in each grid cell of a zooregion (hereafter IndVal and DifVal, respectively). For the IndVal, we used the formula proposed by Dufrêne and Legendre (31) which considers the affinity and fidelity of species to their region. The affinity of the species *i* can be described as *A*_*i*_= *R*_*i*_/*Z*; where *R*_*i*_ depicts the area of the species *i* within its associated zooregion and *Z* depicts the total area of the associated zooregion. The fidelity of the species *i* can be described *as F*_*i*_ = *R*_*i*_/*D*_*i*_, where *D*_*i*_ is the total distribution area of the species *i*. Finally, the IndVal is calculated as *IndVal*_*i*_ = *A*_*i*_ × *F*_*i*_. A species that occupies most of its associated zooregion (i.e. high affinity) and is not present in other zooregions (i.e., high fidelity) will have a high IndVal.

We calculated the DifVal of the non-characteristic species by using a modification of the IndVal, substituting the fidelity by the infidelity (1 − fidelity). Thus, the DifVal was calculated as *DifVal*_*i*_ = *A*_*i*_ × (1 − *F*_*i*_). A species widely distributed within the focal region (i.e., high affinity) but mostly distributed in other zooregions (i.e., high infidelity) will have a high DifVal. Finally, we calculated the robustness value of a grid cell *g* as

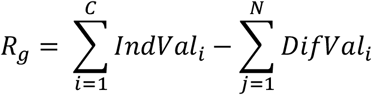

where C represents the characteristic species of the zooregion in the grid cell *g*, and *N* represents the non-characteristic species. Thus, a grid cell containing a large number of highly indicative species and a few species with low diffusion values will have high robustness. We calculated the impacts of introductions and extinctions by measuring how introductions and extinctions alter the sum of IndVal and DifVal values of each grid cell compared to the native scenario. Concretely, we calculated the decrease in the summation of Indval as

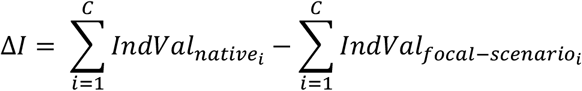

and the increase in the summation of DifVal as

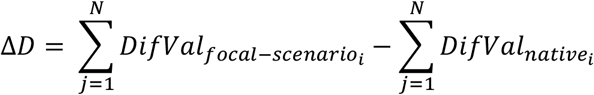

where focal−scenario refers to the introduced or extinct scenario. The total impact of each grid cell is the sum of both values. In that way, a grid cell that loses characteristic species and gains non-characteristic ones would be highly impacted.

## Acknowledgments

We thank the Conservation Biology group of Estación Biológica de Doñana, CSIC, and Elena Marmesat for helpful suggestions. We also thank B. Hawkins, C. Venditti, T. Oliver, O. Gordo, A. Jiménez-Valverde and A. Perrigo for insightful discussions and feedback. This work was supported by the following grants and projects: Agencia Estatal de Investigación from the Ministry of Economy, Industry and Competitiveness, Spain with projects CGL2012-35931 and CGL2017-83045-R AEI/FEDER EU, co-financed with FEDER to E.R., R.B.M., M.G.S., and P.M.L., the Predoctoral Fellowship BES-2013-065753 from the same institution to R.B.M.; a Juan de la Cierva post-doctoral fellowship (JCI-2011-09158) to MG-S; project 707587 H2020-MSCA-IF-2015 from the EU H2020 to M.R. and E.R.; the Swedish Research Council grant 2016-00796 to M.R; and the European Research Council under the European Union’s Seventh Framework Programme (FP/2007-2013, ERC Grant Agreement n. 331024), the Swedish Foundation for Strategic Research, the Swedish Research Council, a Wallenberg Academy Fellowship, the Faculty of Sciences at the University of Gothenburg, and the David Rockefeller Center for Latin American Studies at Harvard University to A.A..

