## Supplementary Materials for "Human activity is altering the world’s zoogeographical regions"

**Appendix A. Community detection algorithm.** The map equation exploits the duality in information theory between compression and detection regularities. In particular, it measures the code length of a modular description of pathways on a network, which in this setting corresponds to sequences of connected grid cells and species. The modular partition of the nodes in the network that gives the shortest description identifies most regularities in the sequences of grid cells and species. In this way, the map equation clusters grid cells with many common species into zooregions. The hierarchical map equation relaxes the constraint that the modular description should only consist of two levels, modules and nodes, and delineates hierarchically nested zooregions.

**Appendix B. Number of delineations.** To assess the zooregional delineation uncertainty, we applied the community detection algorithm multiple times until the solution distribution converged. That is, we assume that we have detected all significant solutions when their relative frequency no longer changes. Specifically, for each scenario, we explored increasingly larger numbers of zooregional clusterings (starting with 100 and increasing by 100) and compared the relative occurrence of different solutions using Fisher's exact tests with p-values estimated from 10,000 Marko Carlo simulations (function fisher.test of the R package stats). We considered that p-values close to one indicated that the solutions had similar relative occurrence, and therefore that we mostly detected all supported solutions. We found that 1500 network clusterings described the zooregional delineation uncertainty for all scenarios and vertebrate groups (SI Appendix, Fig. S10).

**SI Figures.**

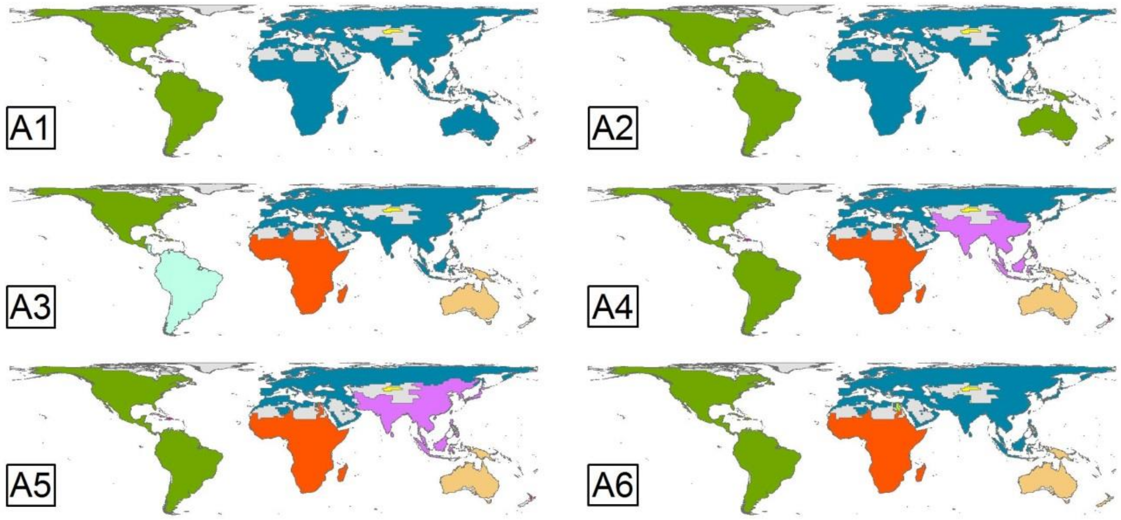


**Fig. S1.** Six solutions for the major zooregions of amphibians. The zooregions represented are the consensus of geographical delineations grouped in each solution. See SI Appendix Table S1 for the detection probability of each solution under each scenario. We called the solutions of the major zooregions of amphibians with the capital letter “A” and a consecutive number.

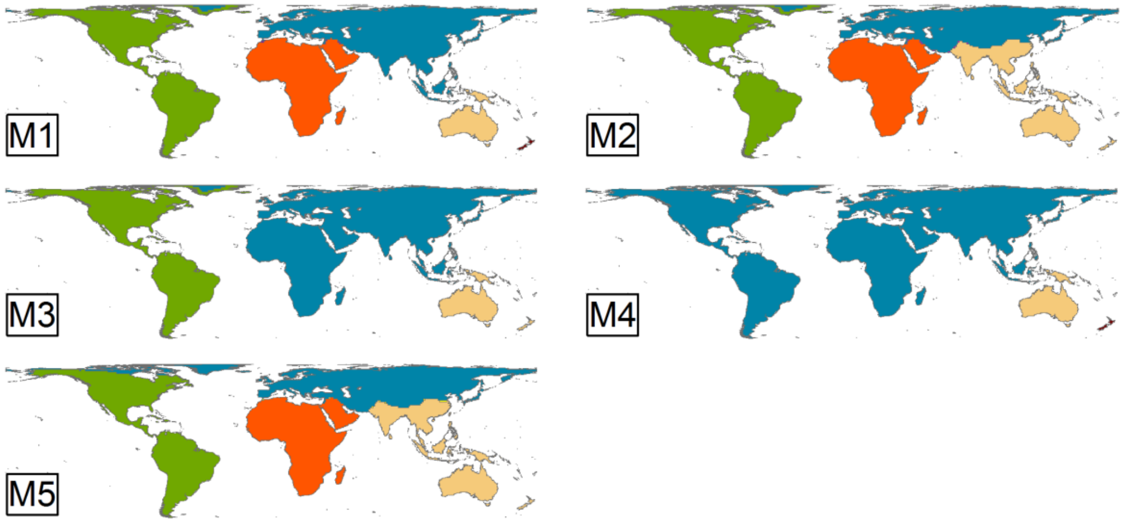


**Fig. S2.** Five solutions for the major zooregions of mammals. The zooregions represented are the consensus of geographical delineations grouped in each solution. See SI Appendix Table S1 for the detection probability of the different solutions. We called the solutions of the major zooregions of mammals with the capital letter “M” and a consecutive number.

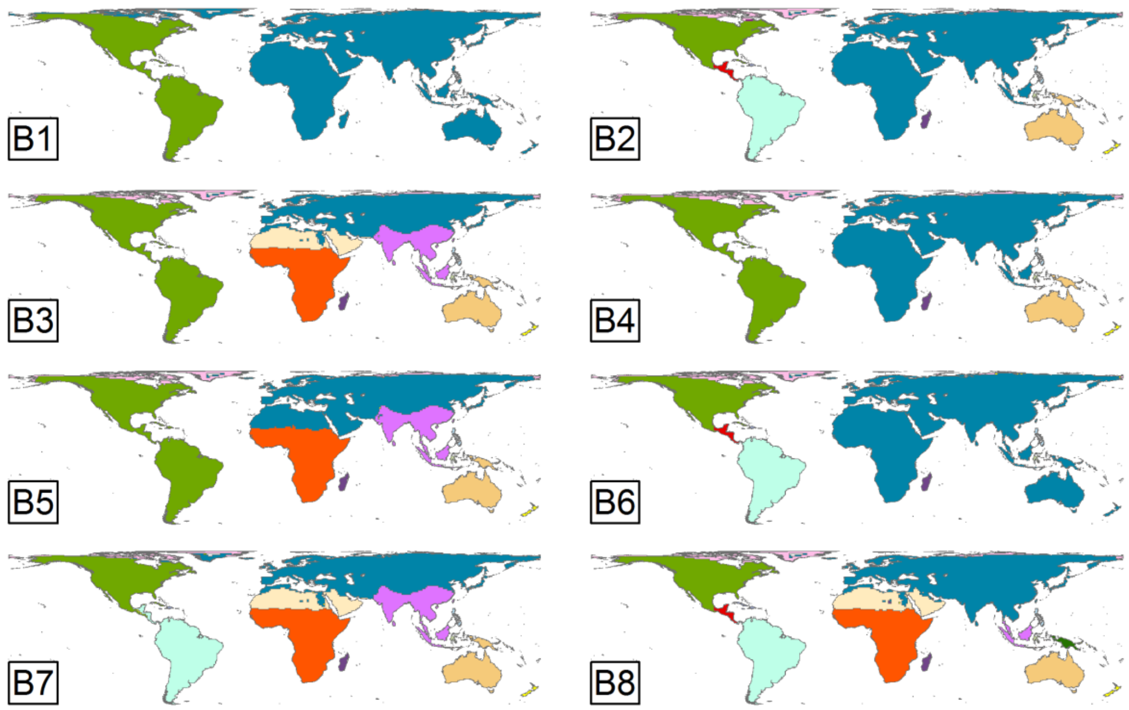


**Fig. S3.** Eight solutions for the major zooregions of birds. The zooregions represented are the consensus of geographical delineations grouped in each solution. See SI Appendix Table S1 for the detection probability of the different solutions. We called the solutions of the major zooregions of birds with the capital letter "B" and a consecutive number.

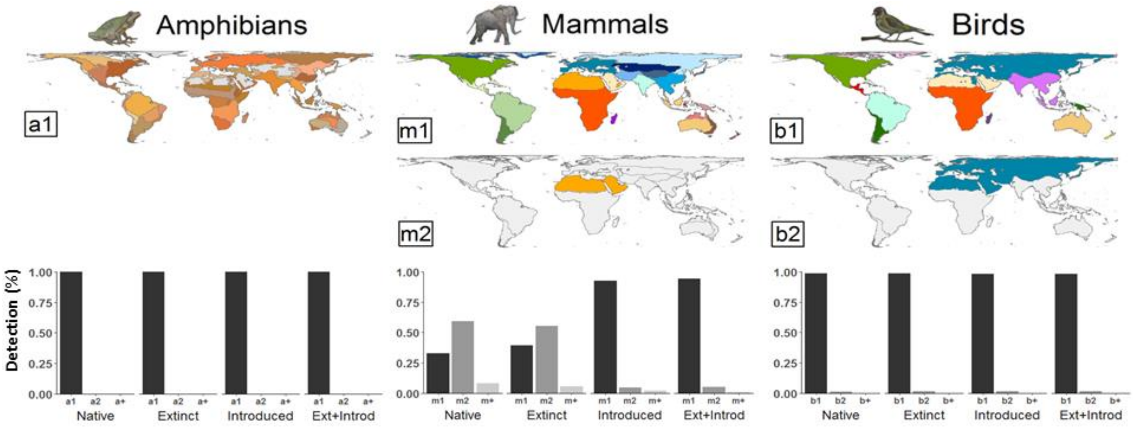


**Fig. S4.** Zoogeographical delineation of the unique zooregion of the minor zooregions of amphibians and zoogeographical delineation of the two most probable solutions for the minor zooregions of mammals, and birds (labeled as a1 for amphibians, m1-m2 for mammals and b1-b2 for birds), and their detectability probability across scenarios (histograms; see SI Appendix Table S1 and Figs. S5-S7 for details). The histograms also show the joint probability of occurrence of supported but non-main solutions, not shown in maps (m+ for solutions 3-11 of mammals; b+ for solution 3 of birds; see SI Appendix, Table S1 for details)

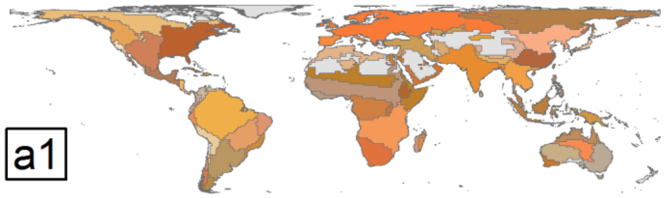


**Fig. S5.** Unique solution for the minor zooregions of amphibians. The zooregions represented are the consensus all delineations. Gray color represent the areas without information about amphibians by the IUCN.

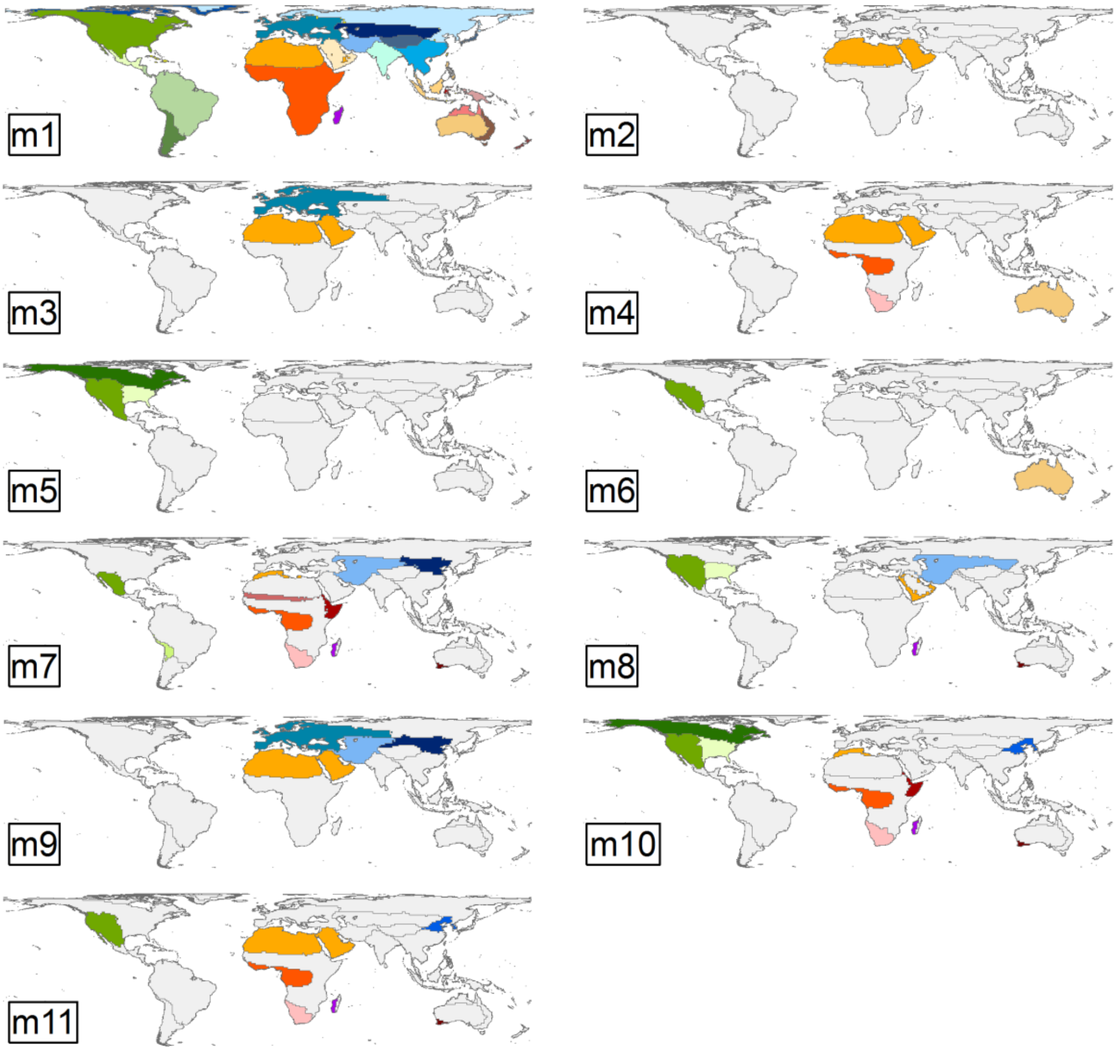


**Fig. S6.** Eleven solutions for the minor zooregions of mammals. The zooregions represented are the consensus of the delineations grouped in each solution. See SI Appendix Table S1 for the detection probability of the different solutions. We called the solutions of the minor zooregions of mammals with the lower letter “m” and a consecutive number. Solution m1 is the most frequent solution in the native scenario and all regions are distinguished by colour. For solutions m2-m11 we used colours only to highlight changes in relationship to m1 (these colours are the same as the general region colour in m1), and used gray colour and line boundaries for zooregions in which no changes were detected compared to m1.

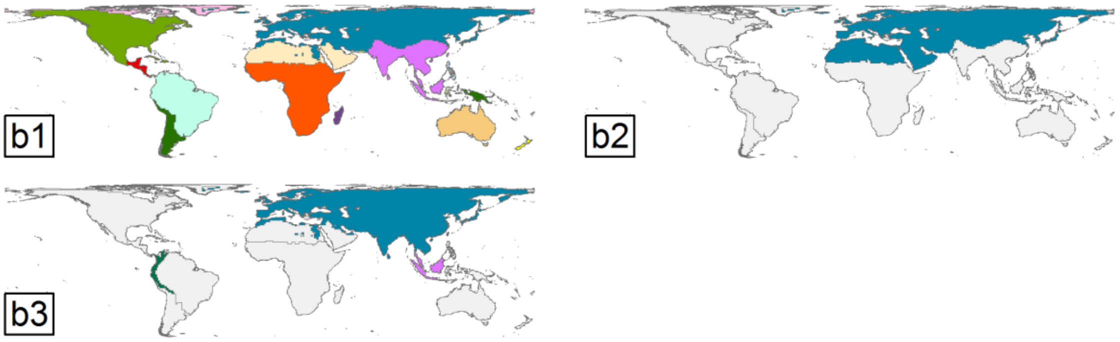


**Fig. S7.** Three solutions for the minor zooregions of birds. The zooregions represented are the consensus of the delineations grouped in each solution. See SI Appendix Table S1 for the detection probability of the different solutions. We called the solutions of the minor zooregions of birds with the lower letter “b” and a consecutive number. The colours of the solutions represent the variations between the solutions with the most frequent solution in the native scenario (b1). The zooregions coloured with gray colour in the solutions b2 and b3 indicate that are like the zooregions of the solution m1 (the most frequent solution in the native scenario). In contrast, the zooregions with non-gray colour in the solutions b2 and b3 are different to the zooregions of the solution b1.

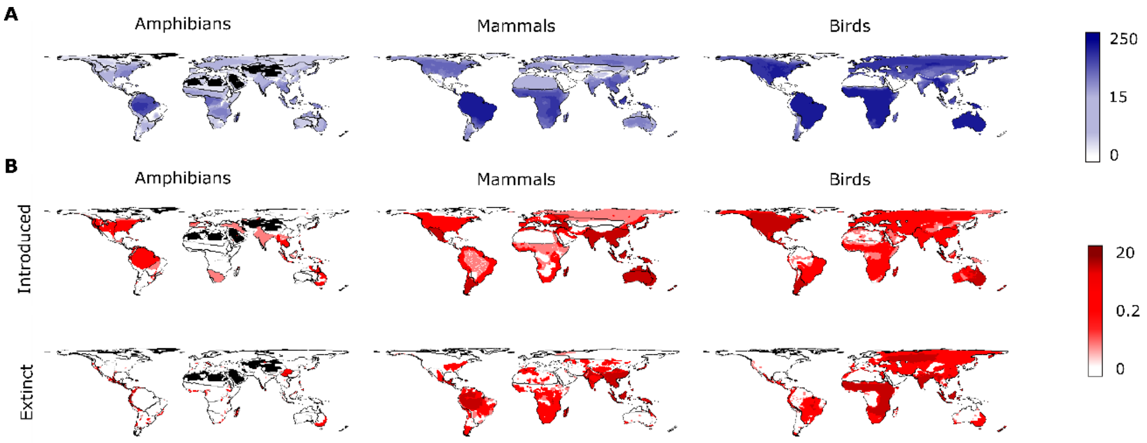


**Fig. S8.** Maps representing robustness and impact for the different scenarios, showing how both vary across regions and vertebrate groups. We represent the (A) robustness and (B) impact values for the grid cells of the minor zooregions, calculated using the most frequent solution for each vertebrate group. (A) Light and dark blue colors indicate low and high relative robustness respectively. (B) Light and dark red colors indicate low and high relative impact respectively.

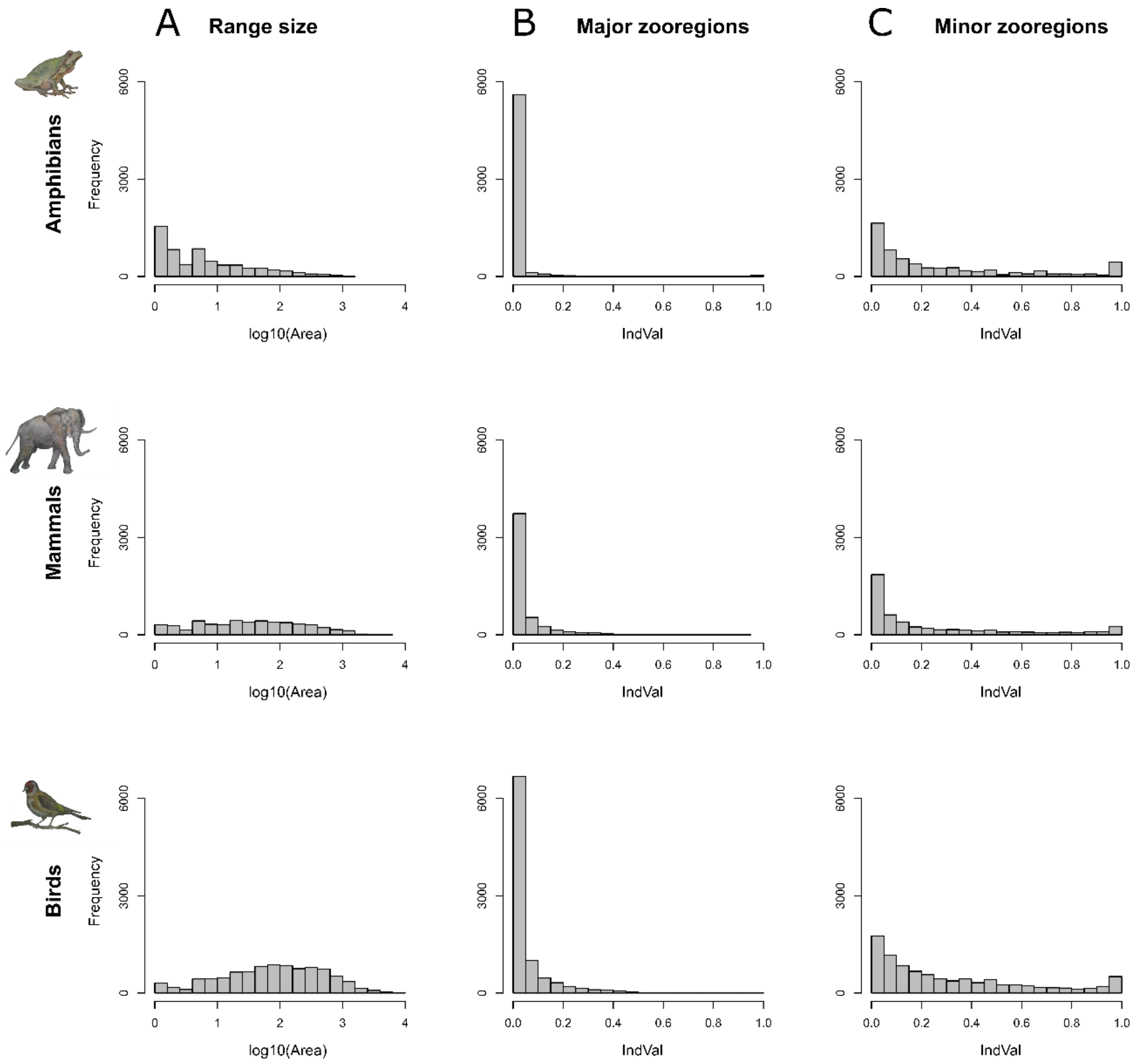


**Fig. S9.** (A) Histograms showing the frequency of species with different distribution areas. The axis x represents the distribution areas of the species in a logarithmic scale for amphibians, mammals, and birds. (B) Histogram showing the IndVal of the species in the major zooregions. (C) Histogram showing the IndVal of the species in the minor zooregions. To calculate the IndVal, we used the major and minor zooregions of the most probable solutions in the native scenario.

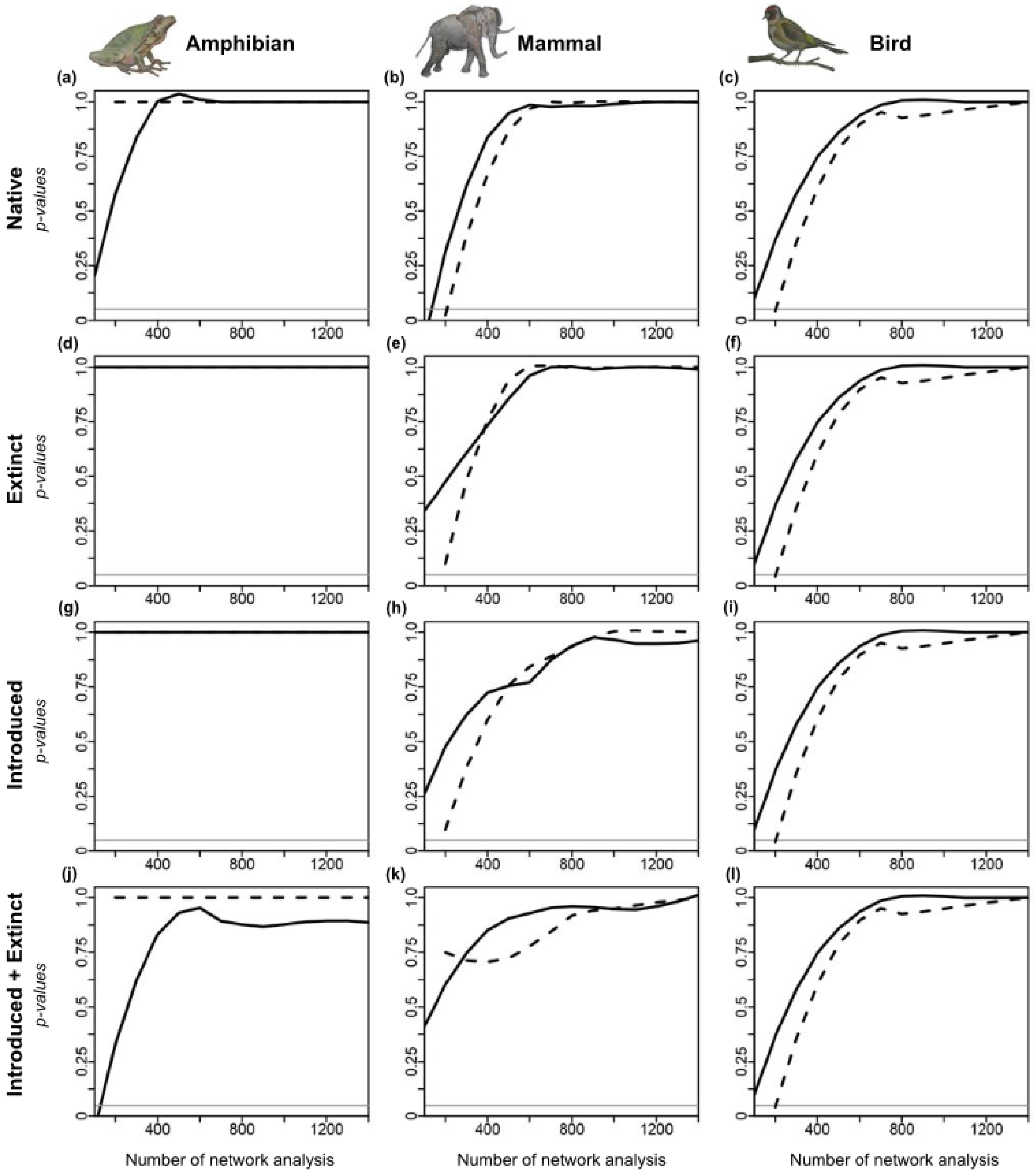


**FS. 10.** Number of community detection analyses needed to represent all alternative supported solutions. We represented the p-values of the Fisher's exact test (y axis) from comparing two frequencies of occurrence for the solutions of the same scenario but using a different number of delineations (x axis). In particular, we compared the frequency of probability of the solutions from a scenario using x number of community detection analyses with the frequency of probability of the solutions from the same scenario using x+100 community detection analyses. Values close to 1 would indicate that frequency of occurrence of the solutions is similar independently of the number of analyses, and that probably new community detection analyses would not provide new supported solutions (see SI Appendix, Appendix B). For our study subjects, we show that 1,500 community detection analyses is enough to identify all alternative supported delineations of zooregions.

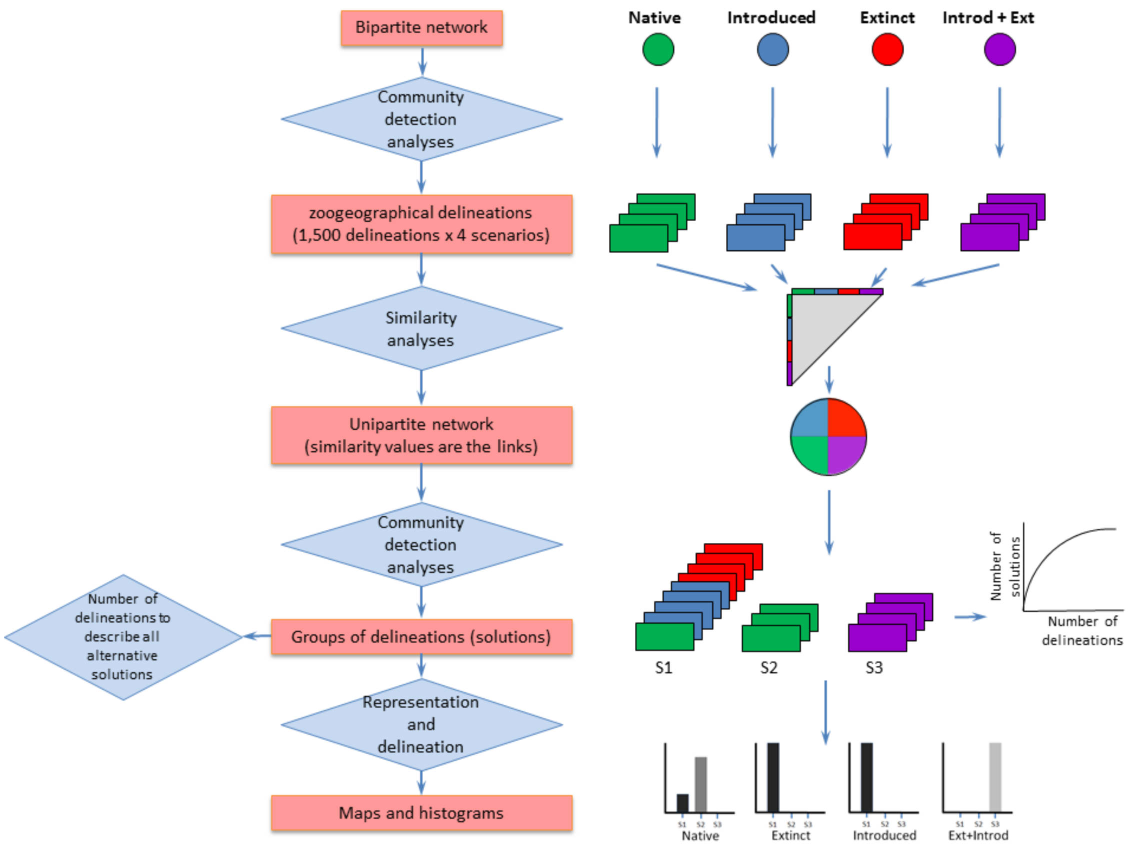


**Fig. S11.** Workflow of the study. The left side of the figure contains the objects (red rectangles) and analyses or activities (blue rectangles). The right side of the figure contains the representation of those objects and analyses.
